# Proteomic analysis of the desmosome identifies novel components required for skin integrity

**DOI:** 10.1101/757203

**Authors:** Kwabena Badu-Nkansah, Terry Lechler

**Affiliations:** Department of Dermatology, Department of Cell Biology, Duke University 27710

## Abstract

Desmosomes are cell-cell adhesions necessary for the maintenance of tissue integrity in the skin and the heart. While the core components of the desmosome have been identified, peripheral components that modulate canonical desmosome functions or that have noncanonical roles remain largely unexplored. Here we used targeted proximity labeling approaches to elaborate the desmosome proteome in epidermal keratinocytes. Quantitative mass spectrometry analysis identified all core desmosome proteins while uncovering a diverse network of new constituents with broad molecular functions. By individually targeting the inner and outer dense plaques, we additionally define proteins enriched in these subcompartments. We validated a number of these novel desmosome-associated proteins and find that many show a dependence upon the core desmosomal protein, desmoplakin, for their localization. We further explored the mechanism of localization and function of two novel desmosome-associated adaptor proteins that we identified, Crk and Crkl. These proteins interacted with Dsg1 and require both Dsg1 and desmoplakin for robust cortical localization. Epidermal deletion of both Crk and CrkL in mice resulted in perinatal lethality with defects in desmosome morphology and keratin organization, thus demonstrating the utility of this dataset in identifying novel proteins required for desmosome-dependent epidermal integrity.

## Introduction

Desmosomes are cell-cell adhesion complexes essential for establishing the mechanical integrity of organs (*1, 2*). They are most abundant in tissues that are regularly subjected to mechanical stress such as the epidermis and cardiac muscle (*3, 4*). Ultrastructurally, desmosomes are highly organized protein complexes with electron-dense plaques whose structural features can be regionalized into nanometer-scale subcompartments (*5*). The extracellular domains of the desmosomal cadherins, desmocollins and desmogleins, form calcium-dependent *trans* dimers with cognate cadherins from neighboring cells. Immediately below the plasma membrane is the outer dense plaque (ODP), where the cytoplasmic domains of these receptors interact with armadillo-domain containing proteins, plakophillins 1-4 (Pkp1,2,3,4) and plakoglobin, which are necessary for efficient formation, clustering, and segregation of desmosome plaques (*6–8*).

Further inward is the inner dense plaque (IDP) where desmosome attachment to intermediate filaments (IFs) is facilitated by the core desmosome protein desmoplakin. Desmoplakin is composed of globular head and tail domains connected by coiled-coils that direct formation of parallel homodimers (*9*). In mature desmosome plaques, the amino terminus of desmoplakin associates with plakoglobin and PKPs in the ODP while the carboxy-terminus anchors cortical IFs in the IDP; therefore, desmoplakin is a bridge that couples desmosome organization to cortical IF attachment (*7, 10, 11*). These core components represent the canonical functional unit of desmosomes necessary for cell-cell adhesion and tethering of IFs to the cell cortex. However, while traditional biochemical fractionation methods (*12–16*), cell reconstitution assays (*7, 17*), and genetic models (*18–23*) were fundamental for functionally establishing this molecular core, other studies have identified additional components that either regulate desmosome adhesion/turnover or mediate non-canonical biological functions at desmosomes. For example, in addition to being stable tethers to IFs, unexpected roles for desmosomes in microtubule (*24–26*) and actin cytoskeleton (*27*) organization have been documented, as well as in signal transduction (*28–31*). It is possible that many of these novel components were not previously identified in early fractionation experiments because they are either peripheral, transient, or biochemically labile components that rapidly dissociate upon cell lysis. Hence biochemically versatile, proximity-dependent approaches, capable of capturing both stable and transiently localized proteins, are necessary to efficiently expand the desmosome proteome.

With the goal of building a more complete parts-list of desmosomes, we turned to proximity proteomic analysis of desmoplakin’s interactome in keratinocytes. Using label-free quantitative MS-based analysis, we identified desmosomal proteins enriched within ODP and IDP regions. We validated the localization of novel desmosome-associated proteins and used mouse genetics to define functional roles for two such proteins, Crk and Crkl. Together, proximity-dependent biotinylation combined with label-free quantitative proteomics broadened the compositional landscape of desmosomes and identified essential new regulators of epidermal integrity.

## Results

### Proximity Biotinylation of Desmosome-Associated Proteins

In order to identify novel desmosome-associated proteins, we turned to BioID for proximity biotinylation proteomics (*32*). During maturation, desmosomes become highly insoluble protein structures. While this has facilitated purification of core desmosomal proteins, the extraction conditions are unsuitable for isolating peripheral and/or transiently associated components.

BioID circumvents this problem by biotinylating proteins localized at desmosomes, thereby making them accessible for purification as they transit the soluble fraction and under harsh conditions. The cytoplasmic face of desmosomes contains two ultrastructurally resolvable units, the inner and outer dense plaques (*33*). In order to target each of these regions we constructed fusions of a mutant biotin ligase, BirA^R118G^ (hereafter called BirA) (*32*) to truncated variants of the core desmosome protein, desmoplakin, which spans these regions (*34*). Fusion to the end of desmoplakin’s head domain (Dsp^N^-BirA) is expected to place BirA in the outer dense plaque while fusion of BirA to the end of the coiled-coil domain (Dsp^CC^-BirA) should place it in the inner dense plaque where desmosomes associate with keratin filaments (Fig. 1A). (*35*). We infected primary mouse keratinocyte cells with lentiviral vectors that enabled stable, doxycycline-inducible expression of each construct along with cytoplasmic-BirA (Cyto-BirA) as a negative control (Fig. 1A). Expression of the fusion proteins was dependent upon doxycycline (Supplemental Figure 1A) and these fusion proteins, tagged with HA, co-localized with endogenous desmoplakin at puncta between cells (Fig. 1B). In addition, desmosomes in Dsp-BirA expressing cells appeared well organized and exhibited stable attachments to keratin fibers (Supp. Fig. 1B). Incorporation of biotin was assayed by staining with fluorophore-conjugated streptavidin, revealing close co-localization of biotin signal with endogenous desmoplakin (Fig. 1C). Line-scan analysis revealed a clear correlation of the biotin signal with endogenous desmoplakin and minimal correlation with E-cadherin and occludin (Fig. 1C and Supp Fig 1C,D). Under conditions where we expressed equivalent levels of BirA fusions, we saw distinct biotinylation patterns by western blot analysis (Fig 1D). Compared with cytoplasmic controls we observed more complex banding patterns in cells expressing either Dsp-BirA fusions.

**Figure 1.**
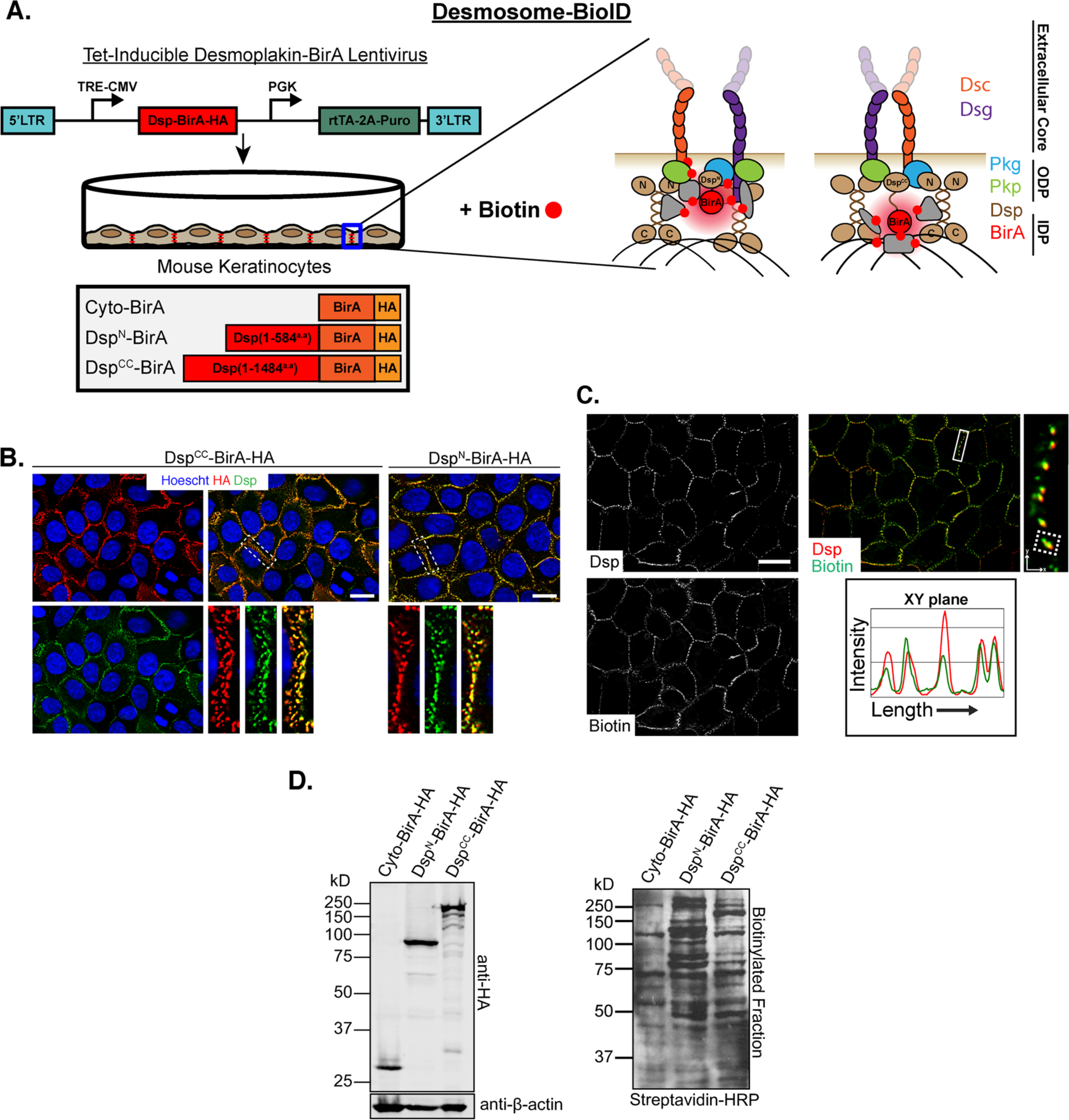
Development of desmosome-BioID in keratinocytes. A) Schematic of doxycyline-inducible BioID approach for targeting desmosomes. BirA, biotin ligase; TRE-CMV, tet-response element promoter; rtTA, reverse tetracycline-controlled transactivator; Dsp, desmoplakin; Dsc, desmocollins; Dsg, desmogleins; Pkg, plakoglobin. B) Validation of Dsp-BirA constructs in mouse keratinocytes immunolabeled for HA tagged-BirA and Dsp. HA, Hemagglutinin fusion tag; Scale Bar, 5μm. C) Immunofluorescent labeling of Dsp^CC^-BirA expressing keratinocytes after overnight incubation in Ca^2+^ and exogenous biotin (50μM). Note that biotinylated proteins labeled with Streptavidin-Alexa488 were enriched at desmosome proximal regions; Scale Bar, 5μm. D) Western blot analysis displays construct expression and biotinylated fraction in biotin-fed, Ca^2+^ induced Dsp-BirA keratinocyte lysates. Membranes were probed with anti-HA and streptavidin-HRP in order to identify Dsp-BirA expression and biotinylation spectrum, respectively.

Interestingly we also observed distinct band patterns between Dsp^N^-BirA and Dsp^CC^-BirA-expressing keratinocytes, suggesting differential enrichment of binding partners. Initial validation of streptavidin precipitates by western blot analysis of Dsp-BioID lysates revealed robust enrichment for representative core desmosome proteins from each subcompartment, with minimal enrichment for adherens junction and tight junction markers, E-cadherin and occludin respectively (Supp. Fig. 1F). Collectively, these results demonstrate Dsp-BioID as a reliable approach for robust labeling and specific purification of desmosome-associated proteins.

### Quantitative Mass Spectrometry Analysis of Desmosome-Associated Proteins

To identify novel desmosome-associated components we performed label-free quantitative mass spectrometry analysis on triplicate samples of biotinylated proteins purified from Dsp^N^-BirA, Dsp^CC^-BirA, and Cyto-BirA (soluble cytoplasmic BirA) expressing keratinocytes. We identified hits as significantly enriched at desmosomes if they passed three constraints: 1) a cutoff of two-fold average intensity over Cyto-BirA, 2) a protein teller probability > 0.8, and 3) a p value of enrichment < 0.01 calculated by student t-test analysis when comparing abundance values of Dsp^N^-BirA or Dsp^CC^-BirA hits to Cyto-BirA controls. By this criteria, we identified 628 proteins enriched in Dsp^N^, and 387 in Dsp^CC^. Of these there were 383 that were enriched in both samples.

We plotted the intensities of significantly enriched proteins for each fusion protein and observed complete coverage of known core desmosome components (Fig 2A). However, our list was not exhaustive, as some previously established desmosome-associated proteins, such as PERP and ninein, were not recovered (*24, 36*). In addition to core desmosome components, Dsp-BioID identified a surprisingly diverse array of proteins from a variety of functional categories (Fig 3A,B). GO term analysis of top hits revealed enrichment of categories related to cytoskeletal and cell-cell adhesion interactions (Fig 2B, Fig 3). Interestingly our list was not limited to regulators of the intermediate filament cytoskeleton, but also included a number of both microtubule and F-actin interacting proteins; consistent with recent work demonstrating that desmosomes can integrate with both of these networks (*25–27, 37*). GO term analysis also revealed novel associations with mRNA and miRNA binding proteins (Fig 2B), an observation previously seen at zonula adherens (*38*), highlighting that desmosomes may share ancillary molecular functions with adherens junctions in addition to their traditional roles in stabilizing cell-cell attachment and cortical cytoskeleton organization.

**Figure 2.**
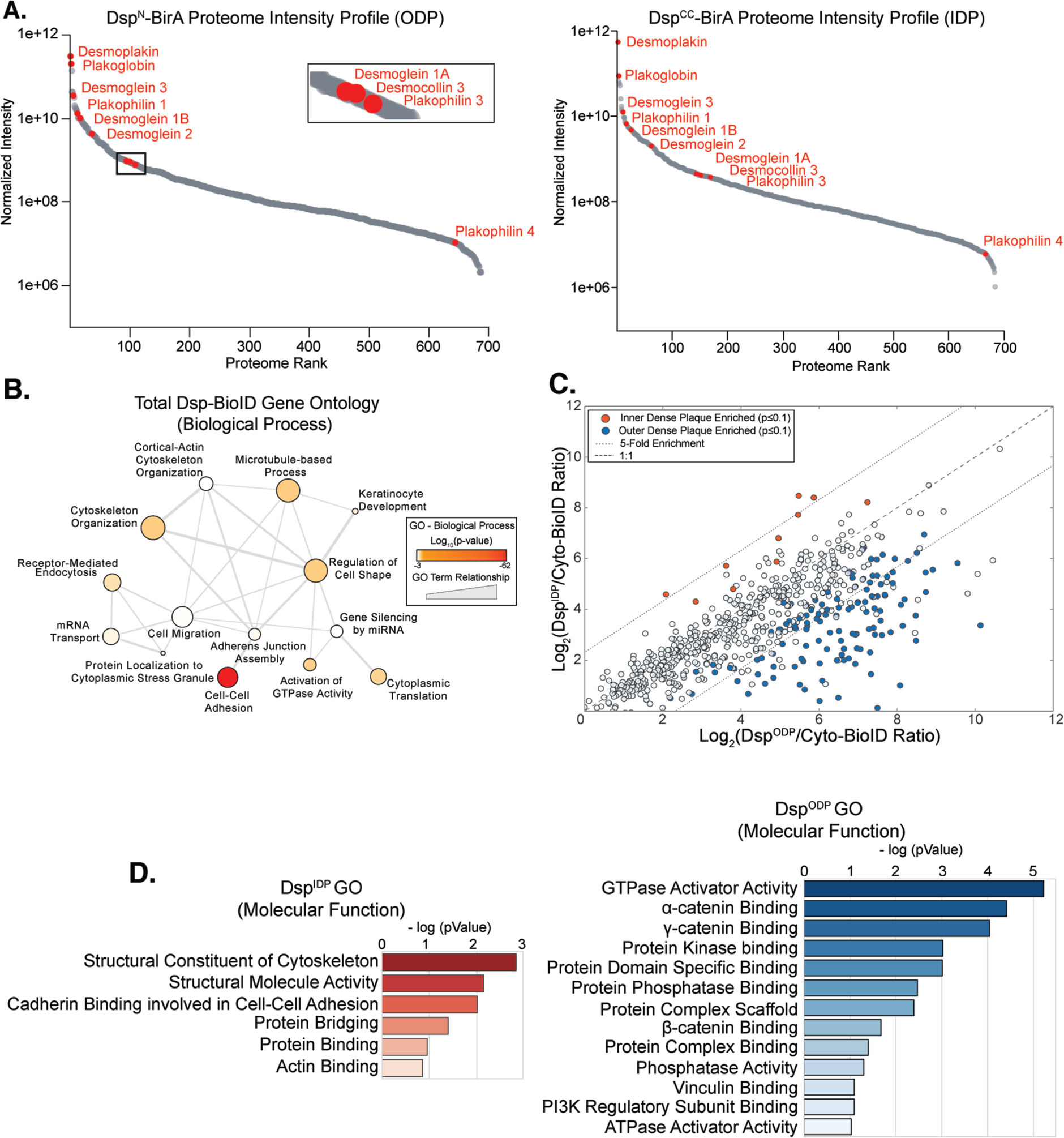
Characterization and quantitative proteomic analysis of desmosome-BioID. A) Ranked intensity plots of protein hits from quantitative proteomic analysis of purified fractions from Dsp^N^-BirA and Dsp^CC^-BirA lysates. Note that in both conditions we observed significant enrichment of core desmosome proteins across large ranges of abundances. ODP, Outer Dense Plaque; IDP, Inner Dense Plaque. B) Revigo plot highlighting functional categorization of biological processes enriched by desmosome BioID. Node size represents relative number of proteins while edge width indicates degree of similarity between connecting GO term nodes. C) Cross correlation plot compares statistical fold enrichment of protein hits from Dsp^IDP^ vs Dsp^ODP^ analyses. Each point represents a protein hit with an associated p-value indicating enrichment in either Dsp^IDP^ or Dsp^ODP^ conditions or neither. D) Graphs of GO terms of molecular funciton for proteins enriched in either Dsp^IDP^ or Dsp^ODP^ conditions.

**Figure 3.**
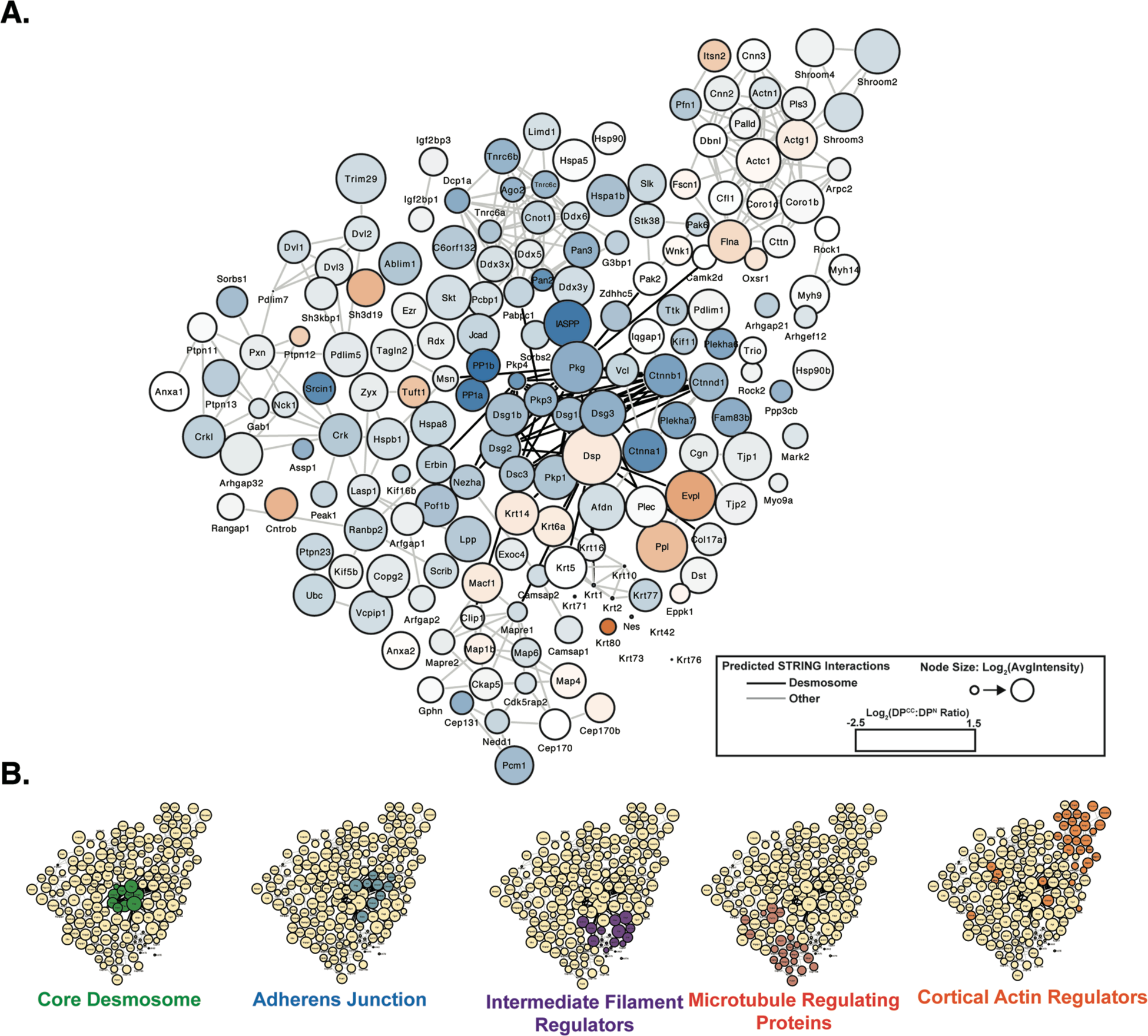
Network graph of the desmosome proteome. A) Top protein hits from desmosome BioID analyses elaborate a rich network of previously known and unknown components at desmosome adhesions. Node titles, sizes, and colors represent gene names; total abundances; and Dsp^CC^ vs Dsp^N^ abundance ratios respectively; edge connections indicate protein interactions identified by STRING; edge color highlights predicted interactions with core desmosome components. B) Network topologies of desmosome BioID protein hits from select functional categories.

Cross-correlation analysis of Dsp^N^-BirA vs Dsp^CC^-BirA hits revealed substantial overlap and positive correlation between both proteomes (Fig. 2C). However, we observed a number of proteins that were statistically significantly more enriched in one compartment versus the other (Fig. 2C and Supplemental Table 1). Protein hits that were more highly enriched by Dsp^CC^-BirA largely included cytoskeletal regulators confirming previous studies and establishing the IDP as a region where desmosome interaction with the cytoskeleton is abundant (Fig 2D). Proteins highly enriched in Dsp^N^-BirA were more diverse in molecular functions and included a number of signal transduction regulators including many phosphatases and signal adaptor proteins (Fig 2D and Supplemental Table 1). These data highlight the ODP as a possible hub for molecular signaling pathways.

### Desmosomes are Essential for Cortical Localization of Diverse Junctional Components

In addition to known desmosome constituents, Dsp-BioID analysis revealed a substantial number of novel desmosome-associated proteins. Intriguingly, we observed that many of these putative components were also identified in previous BioID analyses of adherens junctions (*39, 40*), tight junctions (*41*), or both. Comparative analysis of similar proximity proteomic studies on adherens junctions and tight junctions reveal significant compositional overlap with the desmosome proteome (Supp Fig S2). Since many of these proteins have not been previously shown to localize to desmosomes we speculated that they may exhibit novel cortical localization in cells with high concentrations of desmosomes like keratinocytes. Therefore, we examined the subcellular localization of various high abundant hits using immunofluorescence analysis or fluorescent-protein tagging in calcium-induced mouse keratinocytes. We chose high abundant hits with uncharacterized roles in the epidermis as well as those with known roles but with unspecified localization patterns. Immunofluorescence analysis revealed cortical pools and desmosomal colocalization of Jcad, Ptpn13, Shp2, gephyrin, ExoC4, Pp1-alpha, and Pdlim5 (Fig 4). All of these proteins were also dependent on desmoplakin for their cortical localization. In contrast, Shroom2, tuftelin, and LPP showed only partial co-localization with desmosomal proteins and were either not entirely independent of desmoplakin for this localization, or only moderately affected by its loss (Fig 4H). These data demonstrate that desmosomes have roles in stabilizing the cortical localization of many cortical proteins, including some that they do not closely co-localize with.

**Figure 4.**
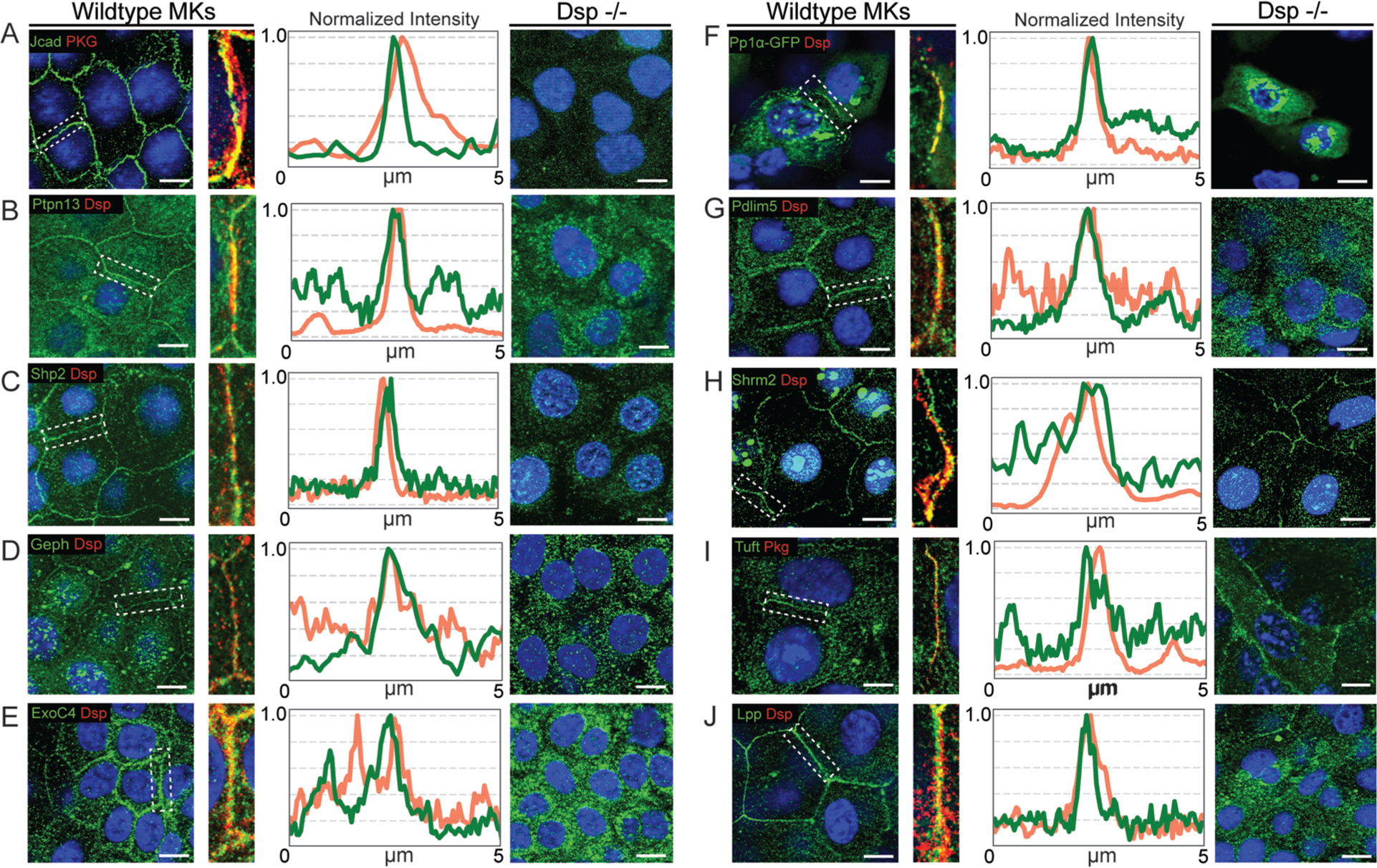
Validation of select desmosome BioID protein targets identify desmosome-dependent and -independent control of cortical complexes. Wildtype, Dsp -/-, or keratinocytes transfected with GFP tagged fusions were fixed and co-stained for desmosomes(red) and proteins identified in our BioID (green). Selected protein hits, listed by gene name, were clustered into groups based on their ability to maintain cortical localization in Dsp null cells. A) through G) represent proteins whose cortical localization were lost in Dsp -/- cells while H) through J) represent proteins whose cortical localization were invariable. Scale bars, 5μm.

### Functional Validation of Crk/Crkl as Desmosomal Regulators

Unexpectedly, amongst soluble signaling components identified by Dsp-BioID we observed an abundance of adapter proteins. We turned to SMART analysis which verified that Src Homology 2 (SH2) and Src Homology 3 (SH3) domains were significantly enriched protein domains in the Dsp-BioID proteome (Fig 5A). We compiled a list of SH3 domain containing desmosome-associated proteins in Supplemental Table 2 and noticed many proteins previously reported to be essential for organizing cortical protein complexes at adherens junctions (*42*) and focal adhesions (*43*). To elucidate further the functional basis behind these observations we focused on Crk and Crkl, paralogous adaptor proteins each composed of one SH2 and two SH3 domains (Fig 5B). Crk and Crkl were identified in both Dsp-BirA proteomes, with both ∼4 times more concentrated in Dsp^N^-BirA over Dsp^CC^-BirA lysates (Fig 5B). Immunofluorescence analysis of endogenous CrkL in mouse keratinocytes revealed punctate cortical localization with robust colocalization with endogenous desmoplakin. A similar localization was seen with a Crk-GFP fusion protein, suggesting that both proteins may have redundant functions at desmosomes (data not shown). While we were unable to observe cortical localization of CrkL in keratinocytes grown in low Ca^2+^, in calcium shift assays, we observed robust desmosome recruitment 3 hours after accumulation of cortical desmoplakin (Fig 5C).

**Figure 5.**
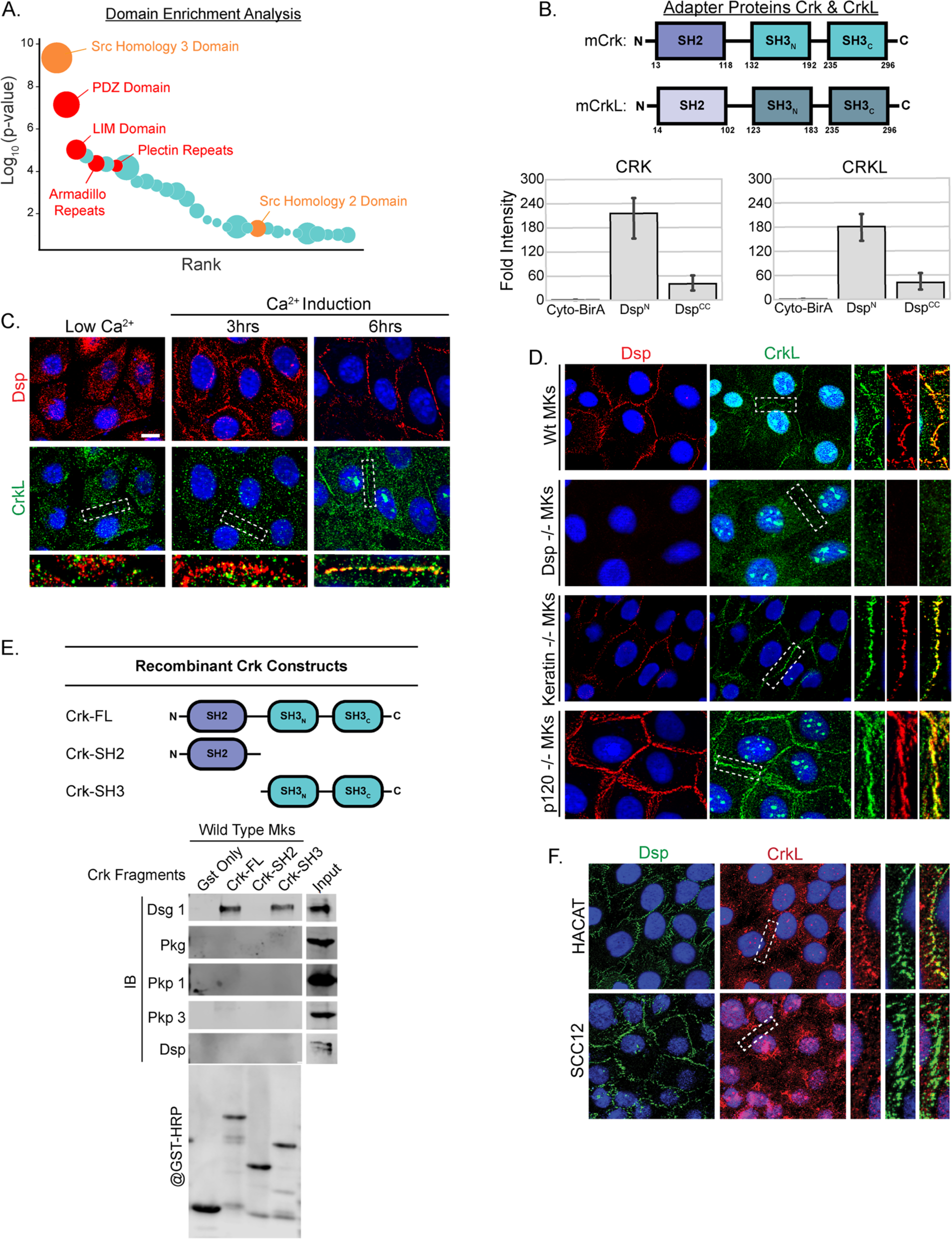
Adapter molecules Crk and CrkL are novel desmosome constituents. A) Domain enrichment analysis of desmosome BioID using SMART. B) *Above:* Schematic representation of domains in Crk and CrkL. *Below:* Graphs displaying fold enrichment of Crk and CrkL in Dsp-BirA proteomes over Cyto-BirA. C) Staining of desmosomes (Dsp, red) and CrkL (green) in mouse keratinocytes before and after Ca^2+^ induction. Note that cortical localization of endogenous CrkL is Ca^2+^ dependent. D) Desmosome (Dsp, red) and CrkL (green) staining in Wt, Dsp -/-, Keratin -/-, and p120- catenin -/- keratinocytes. E) Schematic representation of recombinant full length and truncated GST-Crk constructs. F) Pull down assays from mouse keratinocyte lysates using recombinant Crk constructs reveal associations between Crk and desmoglein 1. G) Desmosome (Dsp, Green) and CrkL (red) staining HACAT and SCC12 keratinocytes. Note cortical loss of CrkL in SCC12 cells also corresponds to decreased Dsg1 expression.

The cortical localization of CrkL was abolished in Dsp-null MKs (Fig 5D). Surprisingly, cortical localization was unaltered in keratin type II null MKs which lack all keratin filaments (Fig 4D). Lastly, cortical loss appeared to be specifically sensitive to desmosome disruption, as cortical levels of CrkL were similar between control and p120-catenin null MKs where adherens junction complexes are largely reduced (Fig 4D) (*44, 45*). These results indicate that cortical recruitment and stabilization of Crk and CrkL in keratinocytes specifically require assembly of mature desmosomes.

To understand interactions that may promote Crk/CrkL recruitment to the cell cortex, we performed pulldown analysis using recombinant, GST-tagged constructs of either full-length Crk or Crk containing either only SH2 or SH3 domains. We found that full-length Crk and Crk^SH3N/C^ each efficiently pulled down Dsg1 from lysates, while the SH2 domain did not (Fig 5E). To gain further evidence for the requirements of this interaction, we examined Crkl localization in HACAT cells, which express Dsg1, and SCC12 cells, a carcinoma line that does not (*46*).

Consistent with Dsg1 promoting cortical localization of Crkl, we observed a lack of desmosomal localization of Crkl in SCC12 cells (Fig 5F). Therefore, Crk/CrkL are desmosome-associated proteins that can physically interact (either directly or indirectly) with core desmosomal components via their SH3 domains. Additionally, we found that Crkl enrichment at the cortex increased upon inhibition of tyrosine phosophatases, suggesting that SH2 domain-phospho-tyrosine interactions may stabilize Crkl at the cell cortex (Supplemental Figure 3).

To determine the physiological roles of Crk and CrkL in vivo, we obtained Crk and CrkL floxed mice (*47*) and mated them to Keratin 14-Cre mice (*48*), allowing for efficient recombination in the epidermis (Fig 6A,B). Homozygous loss of either Crk or CrkL alone did not result in any gross observable phenotype. However, homozygous loss of Crk and CrkL together (Crk/L^dKO^) resulted in neonatal lethality with high penetrance (Fig 6C).

**Figure 6.**
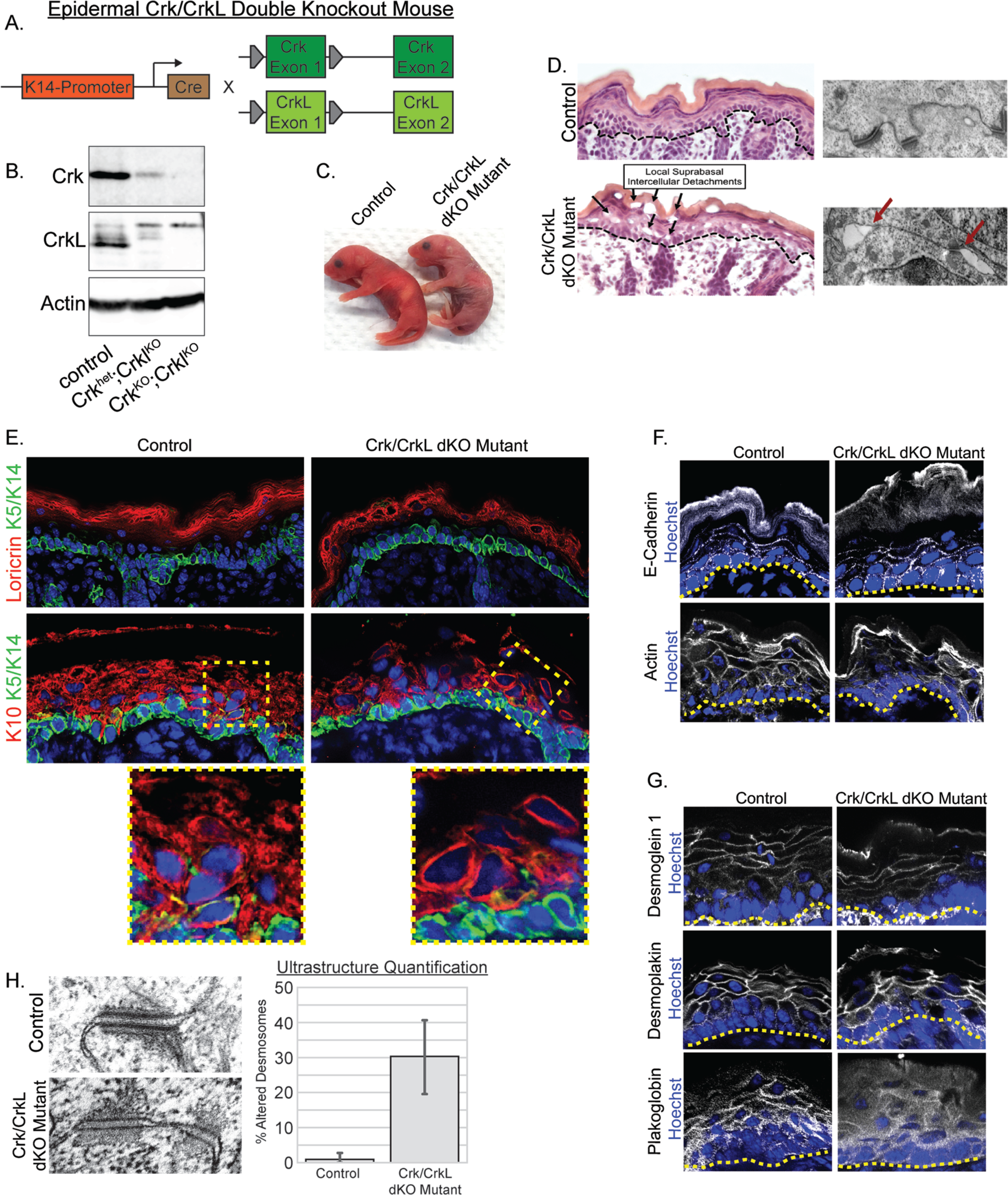
Tissue specific loss of Crk and CrkL results in acantholysis and altered desmosome morphologies in the epidermis. A) Genetic strategy for epidermal targeting of Crk and CrkL. B) Western blot analysis of isolated epidermis from control and dKO mice. Arrow points to band representing loss of CrkL protein. C) Image of control and dKO newborn pups. D) H&E (*left*) and transmission electron microscopy (*right*) images of control and mutant skin. Note intercellular detachments designated by arrows. Scale bars, Left, 30μm; Right, 500nm. E) Immunofluorescent stains of differentiation related markers Loricrin (top, red), keratin 10 (bottom, red), and keratin 5/14 (both, green) in control and dKO epidermis. Inset: Note the perinuclear accumulation of keratin in suprabasal cells of mutant mice. Scale bars, 20μm. F) Expression of e-cadherin (grey) and actin(grey) in control and Crk/CrkL dKO epidermis. Scale Bars, 20μm. G) Immunofluorescent stains of desmoglein 1 (grey), desmoplakin (grey), and plakoglobin (grey) in control and mutant epidermis. Scale bars, 20μm. H) *Left:* High magnification transmission electron micrographs of control and Crk/CrkL dKO epidermis. *Right:* Quantitation of disrupted desmosome organization in the control and Crk/CrkL dKO skin. Scale bars, 100nm.

Histologic examination of newborn epidermis revealed small intercellular separations in the mutant epidermis not found in control mice and consistent with mild desmosome disruption (Fig 6D). Transmission electron microscopy (TEM) of Crk/L^dKO^ skin revealed frequent cell-cell separations throughout the epidermis that were particularly prevalent among differentiated keratinocytes in spinous and granular layers (Fig 6D). Quantitation revealed that desmosome morphology was significantly disrupted in Crk/L^dKO^ skin (Fig 6H). These morphological defects were not associated with increased proliferation or apoptosis (data not shown), nor were there changes in the expression patterns of differentiation markers including keratin 5/14 (basal cell marker), keratin 1 (spinous and granular cell marker), and loricrin (granular cell marker) (Fig 6E).

We next analyzed the localization of desmosomal proteins in Crk/LdKO epidermis by immunofluorescence. While desmoplakin and desmoglein 1 localization appeared largely normal, we observed focal loss of cortical plakoglobin levels scattered throughout mutant epidermis (Fig 6G). While the organization of keratins within the basal layer appeared largely normal, keratin 10 in subrabasal layers was often observed to be collapsed around the nucleus (Fig 6E). In contrast, no defects were noted in either E-cadherin or F-actin localization or levels (Fig 6F). These data demonstrate a physiological role for Crk/Crkl in desmosome stability, keratin organization and epidermal integrity. This provides a proof of principle that the proteomic dataset that we generated increases not only the complexity of the desmosome, but is able to reveal functional regulators of tissue structure.

## Discussion

In this study we used BioID to evaluate the desmosome proteome in epidermal keratinocytes and identify novel desmosome components. Amongst these constituents were proteins with a surprisingly varied array of molecular functions that only partly overlapped with known molecular roles at desmosomes. There were a high number of proteins that harbor adapter domains that stabilize protein interactions necessary for downstream signal transduction. We were able to validate the localization of novel desmosome-associated proteins, and to use mouse genetics to show essential functions in epithelial integrity for the adaptor proteins Crk and Crkl. The function of these adapter proteins, and how they integrate desmosomes with other aspects of cell physiology will be an area of great future interest. Considering that SH2/SH3 adaptor proteins are often involved in tyrosine kinase signaling, the interaction between these networks and desmosomes deserves increased attention. In addition to these novel pathways, our BioID analysis also highlights possible regulators of desmosome stability, such as a single palmitoyltransferase and thioesterase that are excellent candidates to mediate the palmitoylation of desmosomal proteins (*49, 50*).

With this study, proximity proteomics has now been performed for the three major cell-cell junction complexes of vertebrates. Here we show a surprisingly high degree of compositional overlap with BioID analyses of AJs and TJs. While it is likely that some of this overlap is due to labeling promiscuity of BirA, there are two other interesting possibilities for this overlap. First, it may suggest that a number of cortical proteins pass through/near to desmosomes as an essential step of their localization process. An alternative possibility is that desmosomes are permissive for a cortical environment for the localization of various proteins.

It is also plausible that the overlap in AJ and desmosome proteomes may reflect a switch from an AJ-dependent localization in simple epithelia to a desmosome-dependent localization in stratified epithelial, where desmosomes are the predominant cell adhesion structure. Given the extensive similarity in protein families that comprise AJs and desmosomes, it is likely that there is some promiscuity in binding to associated proteins and this may result in functional overlap between these structures in different tissues.

An emerging view is that desmosomes integrate many aspects of cell physiology through noncanonical pathways in addition to their canonical roles in intermediate filament attachment. This work provides hypothesis generating data that will allow future exploration of novel roles for desmosomes in epithelial development and homeostasis.

## Materials and Methods

### Plasmids

Doxycycline-inducible Desmoplakin BirA fusion construct was created by PCR amplifying HA-BirA (Addgene:#36047) and inserting it between Nhe1-Age1 sites in plix402 (Addgene:#41394). Dsp^N^ and Dsp^CC^ truncations were PCR amplified from human cDNA (Addgene#32227) and inserted into plix402-HA-BirA between Nhe1-BstB1 sites. Gephyrin-GFP was a gift from Scott Soderling (Duke University). PP1alpha GFP (Addgene#44224), Crk-GFP(Addgene#50730), CrkL cDNA (Addgene#23354) were all obtained from Addgene. For establishing stable Dsp-BirA cell lines we infected mouse keratinocytes overnight with the relevant lentivirus and subjected cells to puromycin selection. Unless otherwise indicated, mouse keratinocytes were cultured in low Ca^2+^ until time for desmosome induction upon which cell media was switched to high Ca^2+^ media. Keratinocytes were kept in high Ca^2+^ for indicated timepoints up to 48 hours.

### Immunofluorescence

Cells were fixed in either 100% Methanol at -20°C for 3mins or 4% Paraformaldehyde at 4°C for 5mins. Depending on antibody, cells were also pre-extracted in 0.1% Triton at 37C for 30 seconds before fixation. Mouse skin tissue was collected from postnatal day 0 pups, mounted on paper towel, embedded in OCT compound (Sakura 4583), and frozen over dry ice. Frozen blocks were stored long term in -80°C before being cut into 8μm sections on a cryostat and mounted on glass slides. Sections were post-fixed in either 100% Methanol at -20°C for 3 minutes or 4% Paraformaldehyde at 4°C for 7 minutes before permeabilization in 0.1%Triton and subsequent staining. Cells and tissue were blocked and stained in Blocking Buffer (5% Normal Goat Serum, 5% Normal Donkey Serum, 0.1%Triton). Primary antibodies used were CrkL (Santa Cruz, sc-319, sc-365092), Crk (BD Bioscience, 610035), Dsg1 (BD Bioscience, 610273), Desmoplakin (Millipore Sigma, MABT1492), E-Cadherin (Invitrogen, 13-1900), GFP (Abcam, ab13970), HA tag (Roche, 11867423001), JCAD (Santa Cruz, sc-515169), Occludin (Thermo Fisher, 71-1500), Plakoglobin (Santa Cruz, sc-7900; Abcam, ab184919), PTPN13 (Santa Cruz, sc-15356), Shroom2 (GeneTex, GTX100055), Shroom3 (Santa Cruz, sc-376125), Streptavidin-488 (Thermo S11223), and TRITC-Phalloidin (Sigma, P1591). Secondary antibodies used were Donkey Alexa Fluor 488 and 647 conjugated series (Life Technologies/Thermo Fisher) and Donkey Rhodamine Red Conjugated Series (Jackson ImmunoResearch).

### Hematoxylin and Eosin Stain

Frozen tissue sections were prepared as above. Sections were postfixed in 10% PFA at 4°C for 10 minutes. Sections were washed and stained in Mayers Hematoxylin (Sigma-Aldrich, MHS32-1L) solution for 10 minutes, followed by rinsing under running water, 5 dips into Eosin (Polysciences, 09859), dehydration series into 100% ethanol, xylene clearing, and mounting in Permount mounting media. Tissues were coverslipped and sealed using nail polish.

### Mice

All animal work was approved by Duke University’s Institutional Animal Care and Use Committee. Mice were genotyped by PCR and both males and females were analyzed. Mice were maintained in a barrier facility with 12 hour light/dark cycles. Mouse strains used in this study were: Crk fl/fl and CrkL fl/fl (gift from Tom Curran, The Children’s Hospital of Philadelphia) (*47*), and Krt14-Cre (gift from Elaine Fuchs, Rockefeller University (*48*)).

### Microscopy

Cell Stains and tissue sections were imaged on a Zeiss AxioImager Z1 microscope with an Apotome 2 attachment using either Plan-APOCHROMAT 40X/1.3 objective oil objective or Plan-NEOFLUAR 63X/1.4 oil objective, Axiocam 506 mono camera for fluorescent Images or AxioCam MRc camera for H&E Stains, and Zen software (Zeiss).

### Immunoblots

Cells were lysed at 4C in RIPA Buffer (50mM Tris, pH 8, 1% Triton, 150mM NaCl, 0.5% Sodium Dodecyl Sulfate, 50mM Triton, 1 mM EDTA) + Protease Inhibitor Cocktail (Roche, 11697498001), sonicated for 15 seconds, clarified via centrifugation, and stored in -80°C. Lysates were solubilized by mixing 1:1 in loading buffer (10% SDS, 40% Glycerol, 3% Bromophenyl Blue, and 10% Beta-Mercaptoethanol). Proteins in lysates with loading buffer were first denatured by boiling for 5 minutes and cooled on ice for 2 minutes before being loaded into 10% polyacrylamide gels and run for ∼90mins at 125V. Resolved proteins were transferred onto nitrocellulose membranes for 1 hr at 100V. Membranes were then blocked with 5% BSA for an hour before incubation with primary antibody, washed three times in PBST (2.0% Triton in PBS), and the finally incubated with secondary antibodies (Licor, IRDye 680RD Series,CW800 Series). Bands were visualized using a LI-COR Odyssey FC system. Primary antibodies used were Dsg1 (BD Bioscience, 610273), Desmoplakin (Millipore Sigma, MABT1492), E-Cadherin (Invitrogen, 13-1900), Glutathione-S-Transferase (Bethyl Laboratories, A190-122P), HA tag (Roche, 11867423001), Occludin (Thermo Fisher, 71-1500), Plakoglobin (Santa Cruz, sc-7900), Streptavidin-HRP (Thermo Fisher, 43-4323).

### Crk-GST Isolation and Pulldowns

Full length and subdomains of Crk and CrkL cDNA were PCR amplified and cloned upstream of GST between NotI and SalI sites in pGEX-4T1. Recombinant GST fusion proteins were produced in BL21 cells via IPTG induction and isolated from bacterial lysate after sonication in buffer PB (PBS, 1mM EGTA, 1mM EDTA, PMSF, 1mM DTT, Protease Inhibitor Cocktail). Bacterial lysates were clarified through centrifugation and supernatant was incubated in glutathione agarose for 1 hour at 4°C. Crk-GST loaded beads were washed twice in PBST (0.1% Tween-20 +1mM DTT) and fusion proteins were eluted in elution buffer (50mM Tris pH8.0, 5mM reduced glutathione, 1mM DTT). Samples were dialyzed into PBS overnight and protein concentrations were measured via Bradford assay. Dialyzed samples were flash frozen in liquid nitrogen and stored at -80°C.

Crk-GST pulldowns were performed as follows: Recombinant full length and truncated Crk-GST fusions were incubated in clarified lysates from Ca^2+^ induced mouse keratinocytes for 2 hours at 4°C. GST-beads were next incubated with Crk-GST mix for at least 3 hours. Beads were subsequently washed 3 times in wash buffer, resuspended in loading buffer, and eluted via boiling for 5 mins before loading into protein gel for immunoblot analysis.

### Purification of Biotinylated Proteins

Mouse keratinocytes stably expressing Dsp^N^-BirA or Dsp^CC^-BirA were maintained and grown in low Ca^2+^ until 70% confluence. Cells were then incubated overnight with high Ca^2+^(1.5mM) media containing 2μg/ml Doxycycline to induce Dsp-BirA expression concomitantly with desmosome organization. 100μM biotin was next added to the media and incubated for 24hrs before cell lysis (see above). Clarified lysates were incubated with Neutravidin beads (Pierce/Thermo, 29200) overnight at 4°C. Beads were washed once with 2% SDS buffer, once with DOC wash buffer (50mM Hepes pH 7.3, 0.1% deoxycholate, 1% Triton X-100, 500mM NaCl, 1mM EDTA), once with Salt Buffer (10mM Tris pH 8.0, 250mM LiCl, 1mM EDTA, 0.5% deoxycholate, 0.5% NP40), and twice with wash buffer (50mM Tris pH 7.5, 50mM NaCl). For immunoblots beads were resuspended in loading buffer with saturated biotin and for mass spectrometry analysis beads were resuspended in Elution Buffer (250mM Tris pH 6.8, 4% SDS, 0.57M Beta-Mercaptoethanol, 10% Glycerol with Saturated Biotin). In both cases bound proteins were eluted via boiling at 95°C for 5 minutes.

### Mass Spectrometry Analysis

BirA, Dsp^N^-BirA, and Dsp^CC^-BirA samples were each generated and analyzed in triplicate. Each sample were briefly run through a 4-12% gradient SDS-PAGE gel and subjected to in-gel reduction, alkylation, and tryptic digestion. Samples were isolated from gel, lyophilized, and resolubilized in 80μL of 2% acetonitrile/1% TFA supplemented with 10 fmol/μL yeast ADH. From each sample, 5μL was removed to create a QC Pool sample which was run periodically throughout the acquisition period. Quantitative LC/MS/MS was performed on 1μL of each sample, using a nanoAcquity UPLC system (Waters Corp) coupled to a Thermo QExactive Plus high resolution accurate mass tandem mass spectrometer (Thermo) via a nanoelectrospray ionization source. The sample was first trapped on a Symmetry C18 trapping column (5 μl/min at 99.9/0.1 water/acetonitrile), after which analytical separation was performed using a 1.7 μm Acquity BEH130 C18 column (Waters Corp.) with a 90-min linear gradient of 5 to 40% acetonitrile with 0.1% formic acid at a flow rate of 400 nL/min at 55C. Data collection on the QExactive Plus mass spectrometer was performed in a data-dependent acquisition (DDA) mode with a r=70,000 (m/z 200) full MS scan from m/z 375 – 1600 with a target AGC value of 10^6^ ions followed by 10 MS/MS scans at r=17,500 (m/z 200) at a target AGC value of 5^4^ ions. A 20s dynamic exclusion was employed to increase depth of coverage. Following 12 total UPLC-MS/MS analyses, data was imported into Rosetta Elucidator v 4.0 (Rosetta Biosoftware, Inc.), and analyses were aligned based on the accurate mass and retention time of detected ions using PeakTeller algorithm in Elucidator. Relative peptide abundance was calculated based on area-under-the-curve (AUC) of the selected ion chromatograms of aligned features across all runs. The MS/MS data was searched against a custom Swissprot database with Mus musculus taxonomy (circa 2015) with additional proteins, including yeast ADH1, bovine serum albumin, E. coli BirA as well as an equal number of reversed-sequence “decoys” to assess false discovery rate determination. Mascot Distiller and Mascot Server (v 2.5, Matrix Sciences) were utilized to perform the database searches. After individual peptide scoring using the PeptideProphet algorithm in Elucidator, the data was annotated at a 1.3% peptide false discovery rate.

Compiled data was stored in excel files containing raw and normalized quantitative values for each peptide/protein, each replicate, quality control scores, statistical analysis comparisons between Dsp^CC^-BirA, Dsp^N^-BirA, and Cytoplasmic-BirA. Enriched gene lists were imported into DAVID for GO term enrichment analysis (https://david.ncifcrf.gov/summary.jsp). All corresponding charts/graphs were made in Python v3.6.

### Network Analysis

Enriched protein hits in Dsp^CC^-BirA and Dsp^N^-BirA, were determined through statistical comparisons to cytoplasmic-BirA. For both overall desmosome proteome and junctional comparison networks, the top 200 most enriched unique proteins in Dsp^CC^-BirA and Dsp^N^-BirA proteomes were used. Predicted interactions between enriched proteins were determined in STRINGdb (https://string-db.org/). Interaction data, protein symbols with corresponding average intensity readings were imported into Cytoscape v3.4.0 as edges, nodes, and node features respectively, for network visualization. To incorporate proteomes of adherens junctions and tight junctions data for network visualization, data from separate BioID analyses targeting corresponding junctions were used (*39–41*).

## Acknowledgements

We thank Julie Underwood for care of the mice, Tom Curran and Elaine Fuchs for mouse strains, and the Duke Proteomics Core for analysis. This work was funded by the HHMI (KB, Gilliam Fellow) and by NIH grants R01-AR055926 and R01-AR067203 to TL.

---

**Figure S1.**
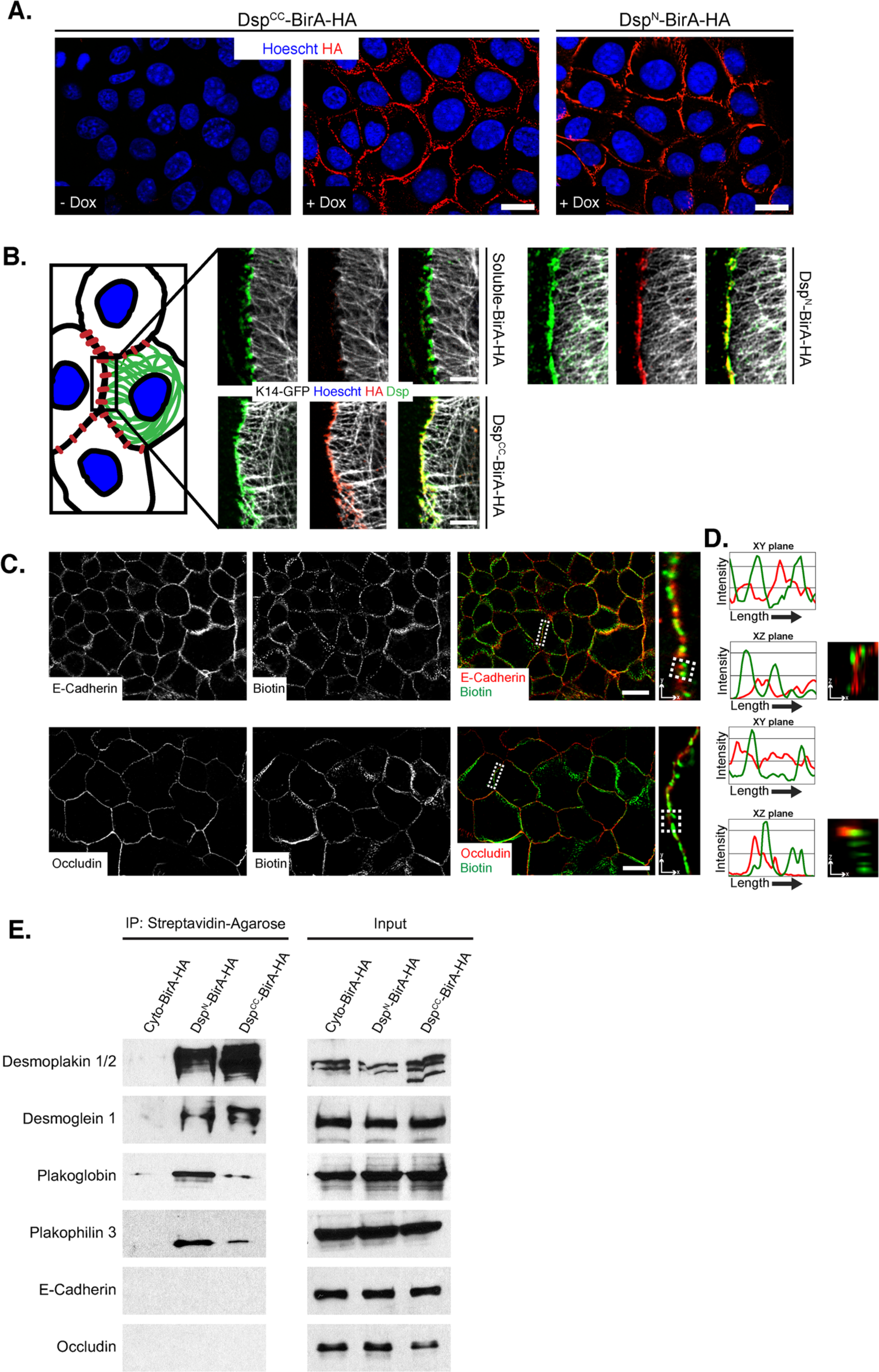
Dsp-BirA fusion efficiently targets desmosomes with high cortical specificity. A) BirA-HA staining (red) of mouse keratinocytes stably integrated with dox-inducible lentivirus of either Dsp^CC^ or Dsp^N^. Scale bar, 5μM B) Dsp-BirA keratinocytes subsequently transfected with K14-GFP and probed for; GFP (grey), HA (Red), Dsp (green). Note that in spite of exogenous expression of Dsp-BirA fusion constructs, desmosomes maintain attachment to keratin network. Scale bar, 3μM. C,D) Staining of biotin (green), E-Cadherin (top red), and Occludin (bottom red) in Dsp^CC^-BirA keratinocytes. Line scan analysis in D) indicate low spatial overlap of biotinylation with e-cadherin (adherens junction marker) and occludin (tight junction marker) stains. Biotinylated proteins were labeled with Streptavidin-Alexa488. Scale bar, 5μM. E) Western blot analysis of junctional components after enrichment of biotinylated fractions from Dsp-BirA lysates.

**Figure S2.**
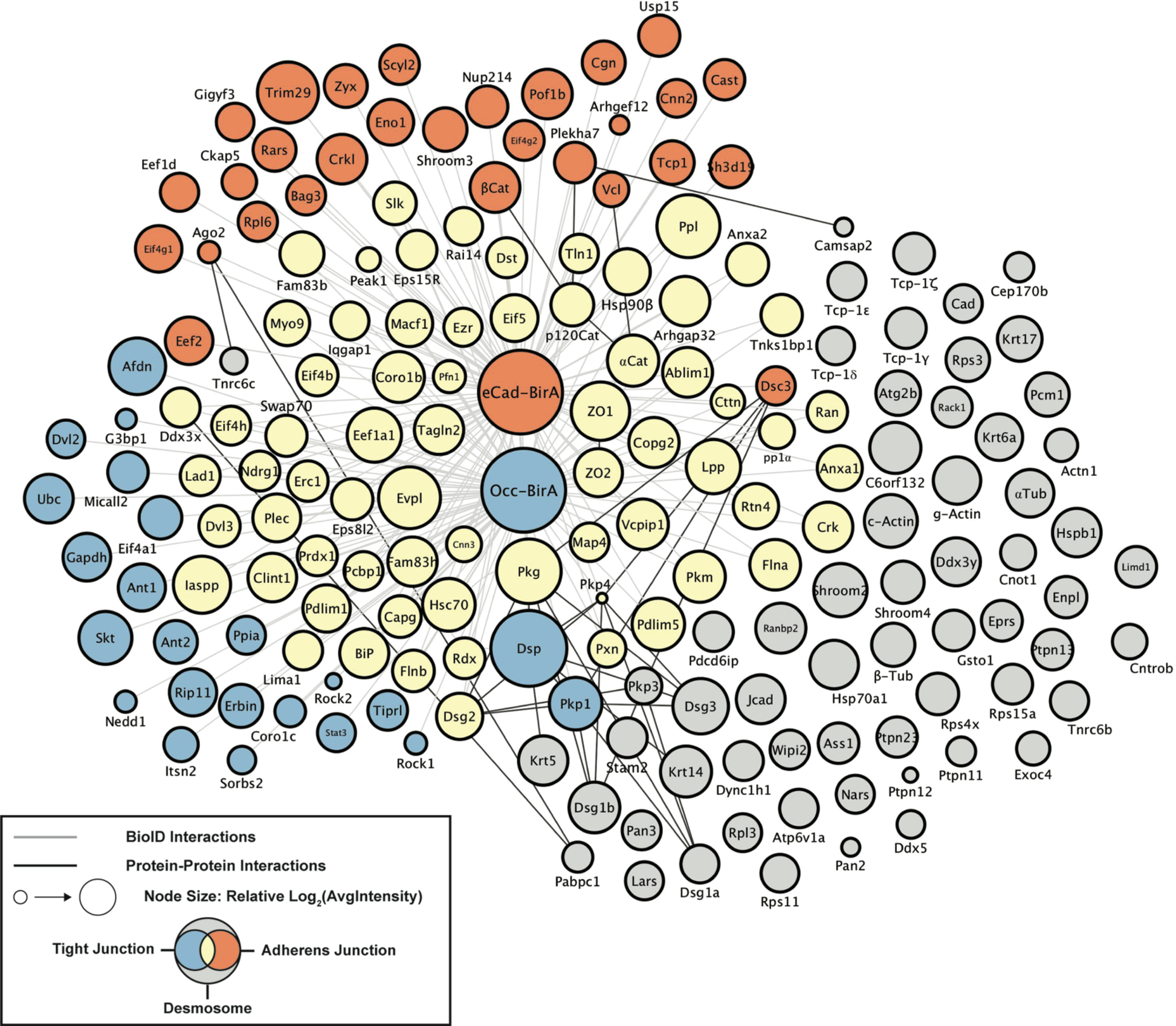
Comparative network analysis of desmosome BioID reveals significant overlap with adherens junction and tight junction proteomes. Comparative analysis of the most abundant desmosome-BioID hits to previously done adherens junction- and tight junction-BioID studies revealed substantial overlap between junctional proteomes. All nodes represent top hits found from desmosome-BioID; node titles represent gene names; node size indicates relative abundances in Desmosome-BioID; node colors represent junctional compartment; and edges represent either BioID interactions (grey) or known protein-protein interactions (black).

**Figure S3.**
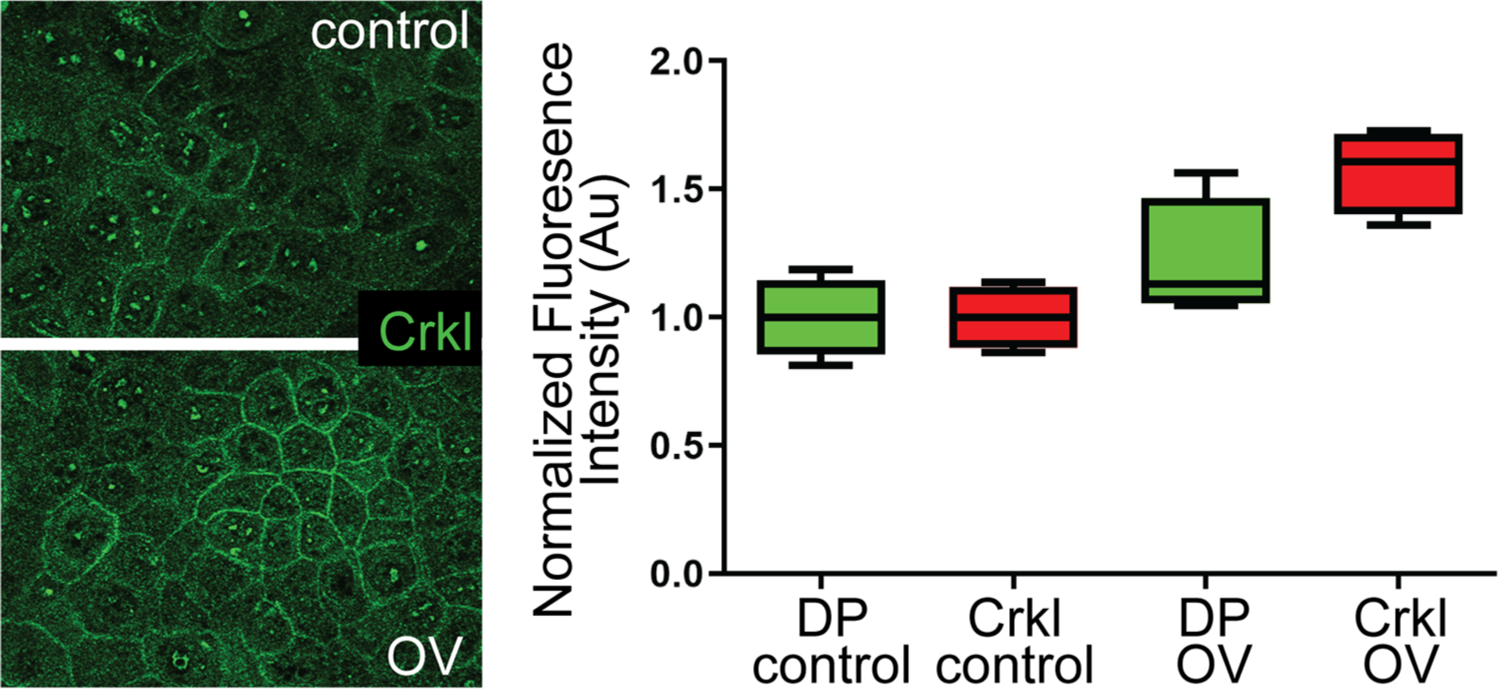
Crkl cortical localization is increased upon inhibition of protein tyrosine phosphatases. Left - Crkl staining in mouse keratinocytes either control treated or treated with orthovanadate for 10 minutes. Right - Quantitation of normalized fluorescence intensity for both desmoplakin (DP) and Crkl.

**Table S1a.**
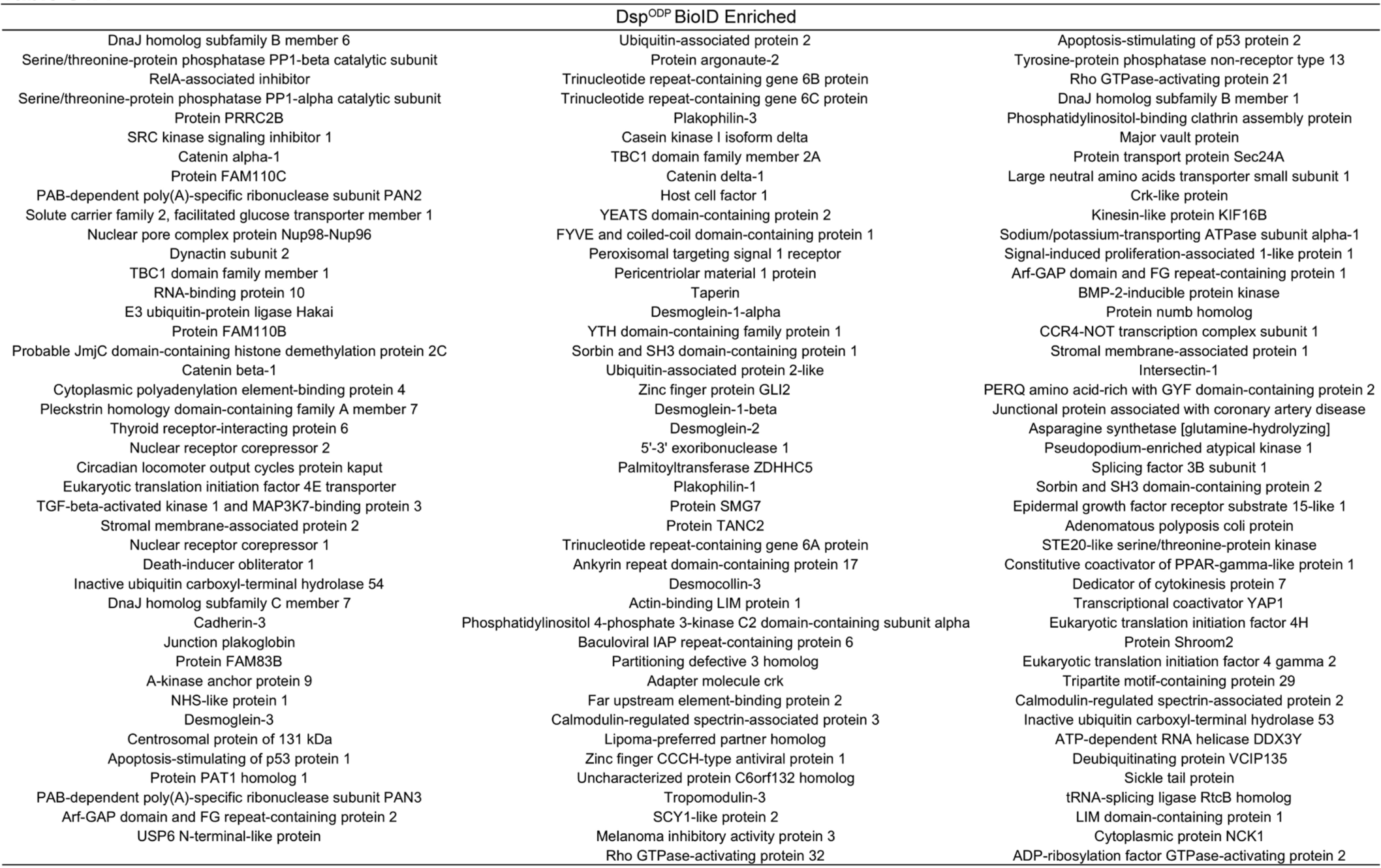

**Table S1b.**
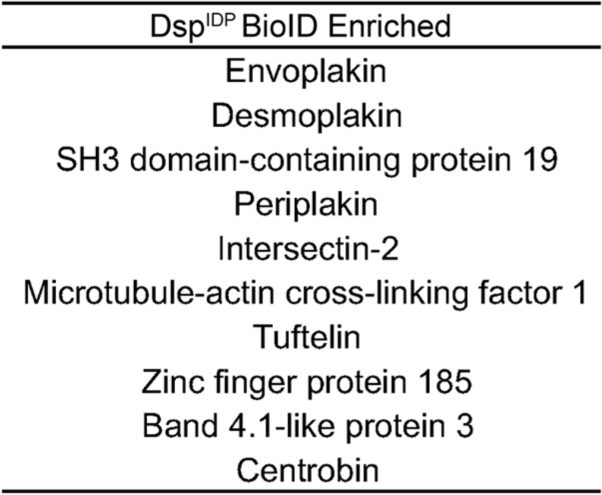

**Table 2.**
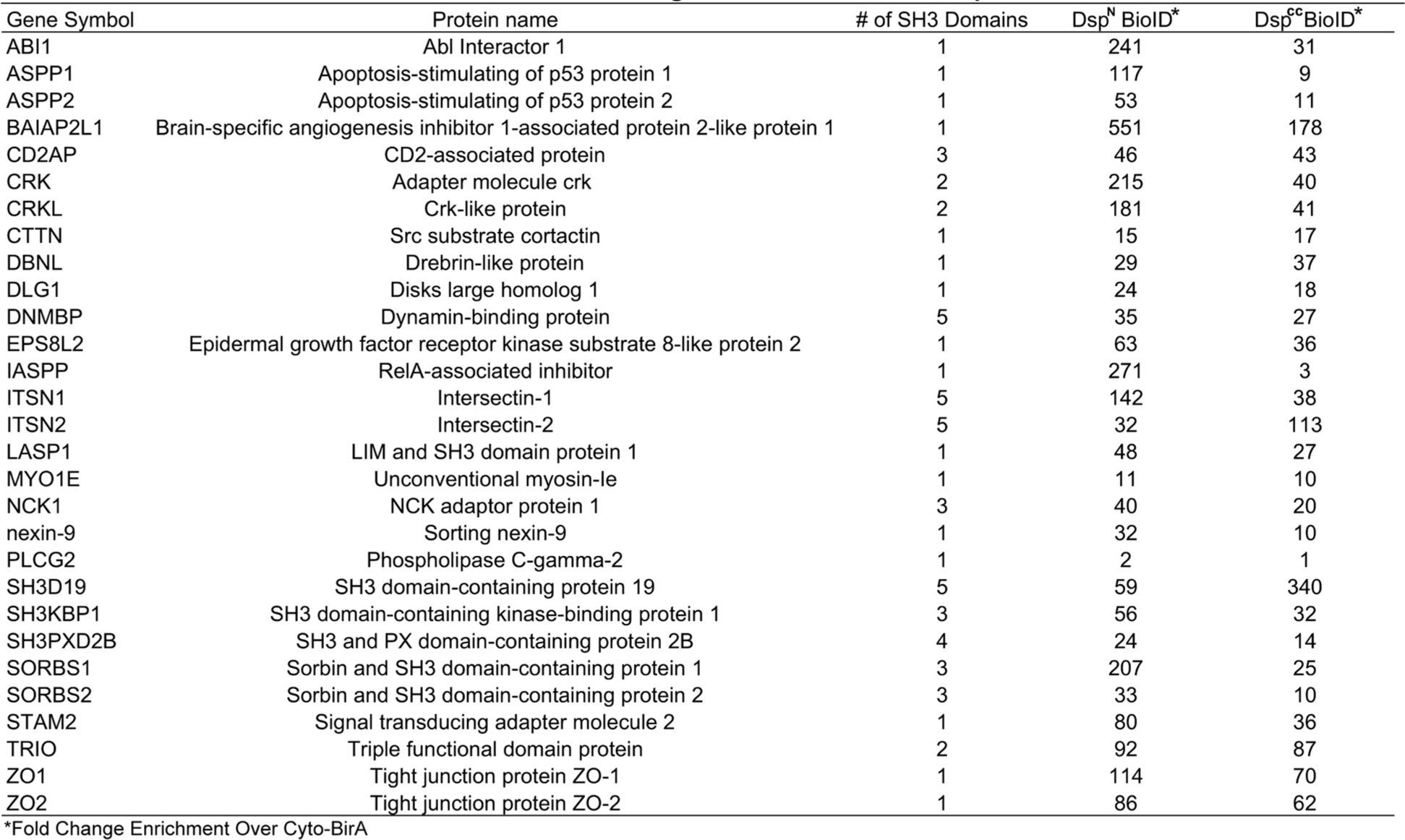
SH3 Domain-Containing Proteins Identified in Dsp-BioID

| Gene Symbol | Protein name | # of SH3 Domains | Dsp<sup>N</sup> BiolD* | Dsp<sup>CC</sup> BiolD* |
| --- | --- | --- | --- | --- |
| ABI1 | Abl Interactor 1 | 1 | 241 | 31 |
| ASPP1 | Apoptosis-stimulating of p53 protein 1 | 1 | 117 | 9 |
| ASPP2 | Apoptosis-stimulating of p53 protein 2 | 1 | 53 | 11 |
| BAIAP2L1 | Brain-specific angiogenesis inhibitor 1-associated protein 2-like protein 1 | 1 | 551 | 178 |
| CD2AP | CD2-associated protein | 3 | 46 | 43 |
| CRK | Adapter molecule crk | 2 | 215 | 40 |
| CRKL | Crk-like protein | 2 | 181 | 41 |
| CTTN | Src substrate cortactin | 1 | 15 | 17 |
| DBNL | Drebrin-like protein | 1 | 29 | 37 |
| DLG1 | Disks large homolog 1 | 1 | 24 | 18 |
| DNMBP | Dynamin-binding protein | 5 | 35 | 27 |
| EPS8L2 | Epidermal growth factor receptor kinase substrate 8-like protein 2 | 1 | 63 | 36 |
| IASPP | RelA-associated inhibitor | 1 | 271 | 3 |
| ITSN1 | Intersectin-1 | 5 | 142 | 38 |
| ITSN2 | Intersectin-2 | 5 | 32 | 113 |
| LASP1 | LIM and SH3 domain protein 1 | 1 | 48 | 27 |
| MYO1E | Unconventional myosin-le | 1 | 11 | 10 |
| NCK1 | NCK adaptor protein 1 | 3 | 40 | 20 |
| nexin-9 | Sorting nexin-9 | 1 | 32 | 10 |
| PLCG2 | Phospholipase C-gamma-2 | 1 | 2 | 1 |
| SH3D19 | SH3 domain-containing protein 19 | 5 | 59 | 340 |
| SH3KBP1 | SH3 domain-containing kinase-binding protein 1 | 3 | 56 | 32 |
| SH3PXD2B | SH3 and PX domain-containing protein 2B | 4 | 24 | 14 |
| SORBS1 | Sorbin and SH3 domain-containing protein 1 | 3 | 207 | 25 |
| SORBS2 | Sorbin and SH3 domain-containing protein 2 | 3 | 33 | 10 |
| STAM2 | Signal transducing adapter molecule 2 | 1 | 80 | 36 |
| TRIO | Triple functional domain protein | 2 | 92 | 87 |
| ZO1 | Tight junction protein ZO-1 | 1 | 114 | 70 |
| ZO2 | Tight junction protein ZO-2 | 1 | 86 | 62 |

\*Fold Change Enrichment Over Cyto-BirA

